# Homogeneity among glyphosate-resistant *Amaranthus palmeri* in geographically distant locations

**DOI:** 10.1101/2020.05.14.095729

**Authors:** William T. Molin, Eric L. Patterson, Christopher A. Saski

**Affiliations:** Crop Production Systems Research Unit, United States Department of Agriculture, Stoneville, Mississippi, United States of America; Department of Plant, Soil, and Microbial Sciences, Michigan State University, East Lansing, Michigan, United States of America; Department of Plant and Environmental Sciences, Clemson University, Clemson, South Carolina, United States of America

**Keywords:** glyphosate, gene amplification, gene homology, evolution, herbicide resistance, *Amaranthus palmeri*, replicon

## Abstract

Since the initial report of glyphosate-resistant (GR) *Amaranthus palmeri* (S) Wats. in 2006, resistant populations have been reported in 28 states. The mechanism of resistance is amplification of a 399-kb extrachromosomal circular DNA, called the *EPSPS* replicon, and is unique to glyphosate-resistant plants. The replicon contains a single copy of the 10-kb 5-enolpyruvylshikimate-3-phosphate synthase (*EPSPS*) gene which causes the concomitant increased expression of EPSP synthase, the target enzyme of glyphosate. It is not known whether the resistance by this amplification mechanism evolved once and then spread across the country or evolved independently in several locations. To compare genomic representation and variation across the *EPSPS* replicon, whole genome shotgun sequencing (WGS) and mapping of sequences from both GR and susceptible (GS) biotypes to the replicon consensus sequence was performed. Sampling of GR biotypes from AZ, KS, GA, MD and DE and GS biotypes from AZ, KS and GA revealed complete contiguity and deep representation with sequences from GR plants, but lack of homogeneity and contiguity with breaks in coverage were observed with sequences from GS biotypes. The high sequence conservation among GR biotypes with very few polymorphisms which were widely distributed across the USA further supports the hypothesis that glyphosate resistance most likely originated from a single population. We show that the replicon from different populations was unique to GR plants and had similar levels of amplification.

## Introduction

In 2006, glyphosate resistant (GR) *A. palmeri* (S. Wats.) was reported in Georgia (GA) [1], and since then, glyphosate-resistant populations have spread to 27 other states as far west as California, and as far north as Maryland (MD) and Wisconsin [2]. The resistance mechanism in *A. palmeri* was identified as amplification and seemingly random insertion of the 5-enolpyruvylshikimate-3-phosphate synthase (*EPSPS*) gene into the genome [3]. *EPSPS* gene sequences from GR and GS plants were identical indicating the resistance was not due to a point mutation. GR biotypes of *A. palmeri* typically contained between 40 and 100 copies of the *EPSPS* gene per genome equivalent [3-7] which increased the C-value of the genome by up to 11% in resistant plants [5]. Interestingly, glyphosate resistance by copy number variation and increased genome size does not seem to lead to immediate consequences to overall fitness [8-10]. The long-term evolutionary consequences of increases in DNA content from targeted amplifications are unknown and may be largely dependent on the level of expression inherited by future generations.

Previous work demonstrated that the amplified *EPSPS* gene was part of a co-amplified 297-kb sequence, the so-called the *EPSPS* cassette [5]. Polymerase chain reaction products from 40 primer pairs distributed equidistantly along this amplified, 297-kb sequence were nearly identical in size and sequence to those in other resistant populations from Arizona (AZ), Kansas (KS), Maryland (MD), Delaware (DE) and Georgia (GA) [6]. However, many of the primer pairs failed to produce products with DNA of GS plants from AZ, KS and GA [6]. These results supported the hypothesis that resistance evolved once, and then spread rapidly by natural means and/or human intervention. However, the study was inconclusive because the primer pairs did not produce overlapping sequences, and therefore were not contiguous, along the 297-kb *EPSPS* cassette [5].

We recently reported that the 297-kb cassette was part of extrachromosomal circular DNA named the eccDNA replicon [11]. The sequence and reference assembly of this eccDNA indicated it was a massive 399,435 bp episome-like DNA containing 59 predicted coding gene sequences, including the *EPSPS* gene, 41 of which were also transcribed [11]. In addition to the *EPSPS*, other genes include multiple AC transposases, replication proteins, a reverse transcriptase, heat shock, and other genes with unknown function [11]. The eccDNA replicon was composed of sophisticated repetitive arrays and was heavily punctuated with sharp changes in A+T and G+C content. It was also found that orthologous genes encoded in the replicon were not found in a colinear organization when aligned to the chromosome-scale reference assembly *A. hypochondriacus* and *A. tuberculatus* [11] which indicates that the eccDNA replicon was assembled from distal parts of the genome instead of a focal amplication of a local genomic segment surrounding the *EPSPS* gene. In a few limited reports, stable, extrachromosomal linear and circular DNAs have been found in plants [12, 13], and the *EPSPS* eccDNA replicon probably represents a novel extension of these. In addition, high-resolution fiber-FISH analysis provided cytological resolution and evidence that the *EPSPS* replicon exists outside the chromosomes in circular forms [14]. In this work, six overlapping BAC tile path probes showed a circular arrangement that includes the *EPSPS* replicon in the GR biotype from MS [14]. In addition to free, extra nuclear circular forms, this eccDNA was also found as chromosomally integrated linear forms (∼30%) and as circular forms attached to metaphase chromosomes through a putative tethering mechanism indicating that the eccDNA is heritable [14]. Unequal segregation of extrachromosomal replicons due to unequal tethering to chromosomes at cell division may account for the nonmendelian inheritance patterns observed in F2 populations [15-19].

Whether the *EPSPS* replicon has single or multiple origins is a pertinent question that impacts both our understanding of resistance evolution, as well as the practical task of GR *A. palmeri* management. A previous study used genotyping-by-sequencing to measure the relatedness of geographically distant, glyphosate resistant populations, however, they were unable to draw a strong conclusion due, in part, to the high amounts of genetic variation in the diploid genome between individuals [20]. *A. palmeri* is dioecious and morphologically diverse, and as such, *EPSPS* replicon sequence diversity may be expected among resistant populations from geographically distant locations, especially if the replicon evolved independently in divergent populations. To further characterize the similarity among the *EPSPS* replicon sequences from divergent GR *A. palmeri* lines and gain evidence for or against a common origin from a single source, a whole genome approach was taken to characterize sequence variation in the entire *EPSPS* replicon as well as genome-wide gene copy number variations in *EPSPS* and other genes in the replicon. In this study, whole genome shotgun sequencing was conducted on GR plants from AZ, GA, KS, DE, and MD and GS plants from AZ, GA and KS, to characterize presence and variation of the eccDNA replicon reference sequence from Mississippi. We show that the *EPSPS* replicon is highly conserved across vast distances in the USA.

## Materials and methods

### Plant sources, resistance confirmation, growth conditions

Seeds were collected from individual GR plants that had survived glyphosate application as previously described [5]. *EPSPS* copy numbers were determined by qPCR on leaves from the third and fourth nodes from two plants from each seedling sample. Copy number assays were performed multiple times as indicated (Table 1). Single populations from other states were provided by M. Jugulam, Kansas State University, W. McCloskey, University of Arizona, B. Hoagland, USDA-ARS, and M. VanGessel, University of Delaware.

**Table 1.** EPSPS copy numbers for the eight biotypes used herein by qPCR compared to the copy number variation (CNV) determined by the CNVnator program.

| Origin | Glyphosate sensitivity | Average Copy Number | Number of DNA isolations per seed source | CNVnator estimate of read depth |
| --- | --- | --- | --- | --- |
| Arizona | Resistant | 77 | 4 | 279 |
| Arizona | Sensitive | 1 | 4 | 0.4 |
| Georgia | Resistant | 35 | 2 | 177 |
| Georgia | Sensitive | 1 | 2 | 2.5 |
| Kansas | Resistant | 69 | 4 | 156 |
| Kansas | Sensitive | 2 | 4 | 2.8 |
| Maryland | Resistant | 35 | 3 | 271 |
| Delaware | Resistant | 38 | 3 | 238 |
| Mississippi-13 | Resistant | 38 | 12 | -- |

Plants were grown in 9 × 9 × 9 cm plastic pots that contained a commercial potting mix (Metro-Mix 360; Sun Gro Horticulture, Bellevue, WA, USA). Seeds were sown on the potting mix surface and lightly covered with 2 mm of potting mix. Pots were sub-irrigated and maintained in a greenhouse set at a temperature regime of 30/25 °C (day/night) and a 15-h photoperiod under natural sunlight conditions supplemented with high-pressure sodium lights providing 400 µmol m^−2^ s^−1^. Sampling for whole genome sequencing was performed using a leaf from the third node of two representative plants from each population.

### qPCR determination of *EPSPS* copy number

qPCR was performed to determine copy numbers of *EPSPS* in plants from the eight plant sources using primer pairs CAACAGTTGAGGAAGGATCTG (AW146) and CAGCAAGAGGAAGGATCTG (AW147) [6]. Acetolactate synthase gene (*ALS*) was used as a housing keeping gene with primers GCTGCTGAAGGCCTACGCT (AW23) and GCGGGACTGAGTCAAGAAGTG (AW24) [6]. DNA was extracted from two leaves each from different seedlings. Reactions consisted of 10 ng of genomic DNA,1.5 μM primers, 1 X Power Sybr Green master mix (Life Technologies), and H2O to 50 μL. Reactions were performed using an ABI 7500 Real Time PCR System. Cycle conditions are as follows: 50 °C for 2 min, 95 °C for 10 min, 40 cycles of 95 °C for 15 s and 60 °C for 1 min, and a 4 °C hold. A melt curve analysis was included. Data were analyzed according to the standard curve method and error was calculated.

### Whole Genome Shotgun Re-sequencing and Variant Discovery

Whole genome shotgun sequencing of 5 GR (AZ-R, KS-R, GA-R, DE-R, MD-R) and 3 GS (KS-S, GA-S, AZ-S) biotypes was performed on total genomic DNA isolated by standard CTAB plant DNA extraction procedures as previously described [6]. DNA was extracted from two leaves each from different seedlings. Sequencing libraries were prepared by ultrasonic shearing of the gDNA with a Covaris ultrasonicator (Covaris) to an average size of 500-bp and indexing with sequencing adapters using the TruSeq DNA (PCR free) kit (Illumina). Sequencing was performed on a single lane of an Illumina HiSeq2500 with 500 cycles in high-output mode. Raw data was preprocessed to remove adapter and low-quality sequences with the Trimmomatic software [21], and cleaned reads mapped to the reference *EPSPS* replicon sequence with the Bowtie2 short read mapper using default parameters [22]. BAM alignment files were filtered for concordant read alignments and mapping quality with the Samtools toolset (v1.3.1) and read depths were determined with the Samtools ‘depth’ function [23]. Single nucleotide variants were determined with the HaplotypeCaller walker of the GATK v3.5 software package and outputted in variant call format (VCF) [24]. Variant sites were filtered for depth (DP>10) and mapping quality (MQ>30) with vcftools [24]. Circular figures were plotted with the Circos plotting tool [25]. Cleaned reads were aligned to the entire AZ-S genome assembly modified by concatenating with the entire replicon sequence using Bowtie2 short read mapper [22]. BAM alignment files were filtered for concordant read alignments and mapping quality with the Samtools toolset [23]. These BAM files were then analyzed for copy number variants (CNV) using CNVnator [26, 27] using standard parameters and a bin size of 500bp. The outputs were then turned into BED files and intersected with the annotation BED file for the whole genome using BedTools [28] to call copy number variants for every gene in the genome and the replicon.

### *Amaranthus palmeri* (GS) genome, *EPSPS* reference sequence, and WGS availability

The *Amaranthus palmer* (GS) reference sequence for this article can be found in the EMBL/GenBank data under BioProject ID PRJNA606296; Submission GenBank SAMN14120689. The *EPSPS* reference sequence (GR) for this article can be found in the EMBL/GenBank data under BioProject ID PRJNA413471; Submission GenBank MT025716. Whole genome resequencing datasets for each state and biotype used in this study can be found in the EMBL/GenBank data under BioProject ID PRJNA413471; Submission Genbank SAMN13521421-SAMN13521428.

## Results and discussion

The *EPSPS* gene copy numbers were determined for the five GR biotypes and three GS biotypes and are shown in Table 1. The standardized read depth for every gene in the eight *A. palmeri* whole genome resequencing datasets was calculated by CNVnator with a sliding 500bp window and is presented in Supplemental Table 1. Relative copy number of EPSPS determined by qPCR was proportional with relative read-depth of the replicon as determined by the software CNVnator.

In order to analyze *EPSPS* replicon representation, variation, and configuration, sequence reads were aligned as pairs to the EPSPS reference, which contains a single copy of the *EPSPS* gene flanked by 135-kb of sequence upstream and 244-kb downstream (Fig. 1). The outer blue ring represents the consensus sequence of the 399,435 bp contig. Consecutively numbered gene annotations are presented as histograms in the light gray inner track (*EPSPS* is PA25) and the direction of transcription of putative transcribed genes is indicated by the color exterior and interior of the center axis of the light gray band, eg. green and exterior is clockwise and yellow and interior is counterclockwise (Fig. 1A). The next two broad, concentric inner rings, gray and blue, represent the results of WGS sequencing and mapping of read pairs from GR and GS samples from Mississippi to the *EPSPS* consensus sequence. This approach provided a means to assess representation, contiguity, and variation between the sensitive and resistant biotypes (Fig. 1A). The gray and blue concentric rings are graphic representations of read depths up to ∼8,000 reads covering certain bases. The light orange glyph (along outer edge of gray circle) against the gray background depicts read depths from GR plants. Dark orange indicates bases that are covered by greater than 2,000 reads. The glyphs along the outer edge of the blue track are incomplete indicating gaps in sequence for reads from sensitive plants. The read depth in GR confirmed the presence of a contiguous circular replicon sequence and the lack of contiguous sequence in GS (Fig.1A). The red/orange spikes in both tracks are areas of higher read depth represent sequences common to both genomes, possibly originating from low-complexity sequence, repetitive elements, or other high copy number DNAs present in higher abundance in the genome (Fig. 1A and 1B). These spikes may indicate that the replicon is a re-arranged assembly of the GS genome because of the lack of concordant read pair alignments from the GS WGS datasets.

**Figure 1A.**
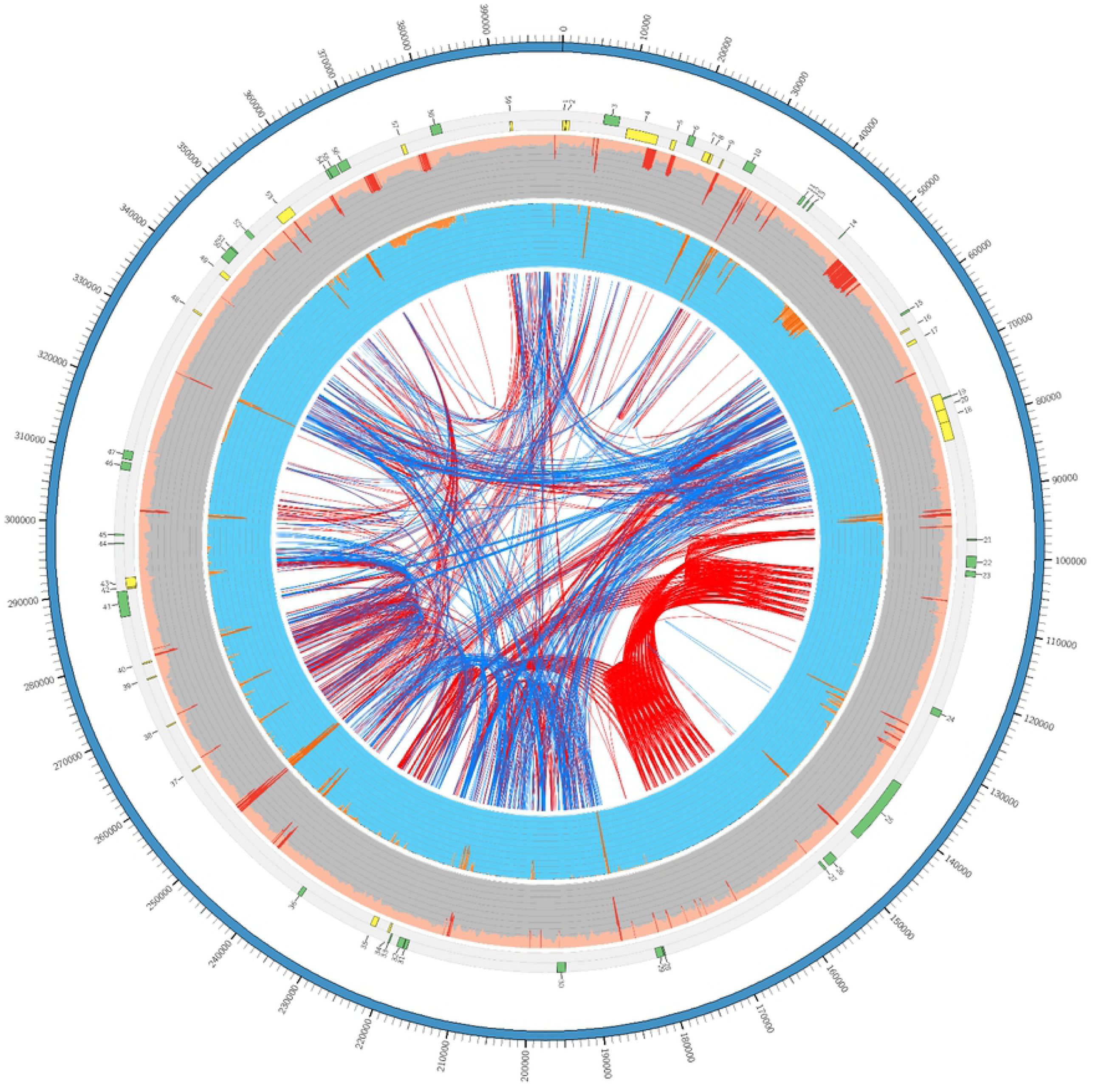
The 399,435 kb *EPSPS* replicon. Whole genome shotgun resequencing of glyphosate resistant and susceptible biotypes and mapping of reads of both biotypes (GR and GS) to the reference *EPSPS* replicon was performed. The outer ideogram (blue) is the 399, 435 kb reference assembly. The outer gray and inner bright blue tracks are circular graphs, the diameter of which represents 12000 WGS reads. The light pink band along the outer edge of the gray track is the read depth for a GR plant and indicates a read depth about 2000 whereas a similar continuous track is not present from GS plants. The inner bright blue track represents GS WGS coverage. In the gray background, light pink (GR) colors, and dark red mark regions covered by greater than 2,001 reads. Read depths for GS biotypes are orange. Interestingly, WGS depths for GR were congruent in depth across the *EPSPS* replicon; while the GS WGS data contained gaps. Tandem repeats are indicated in red and indirect repeats in blue.

**Fig. 1B.**
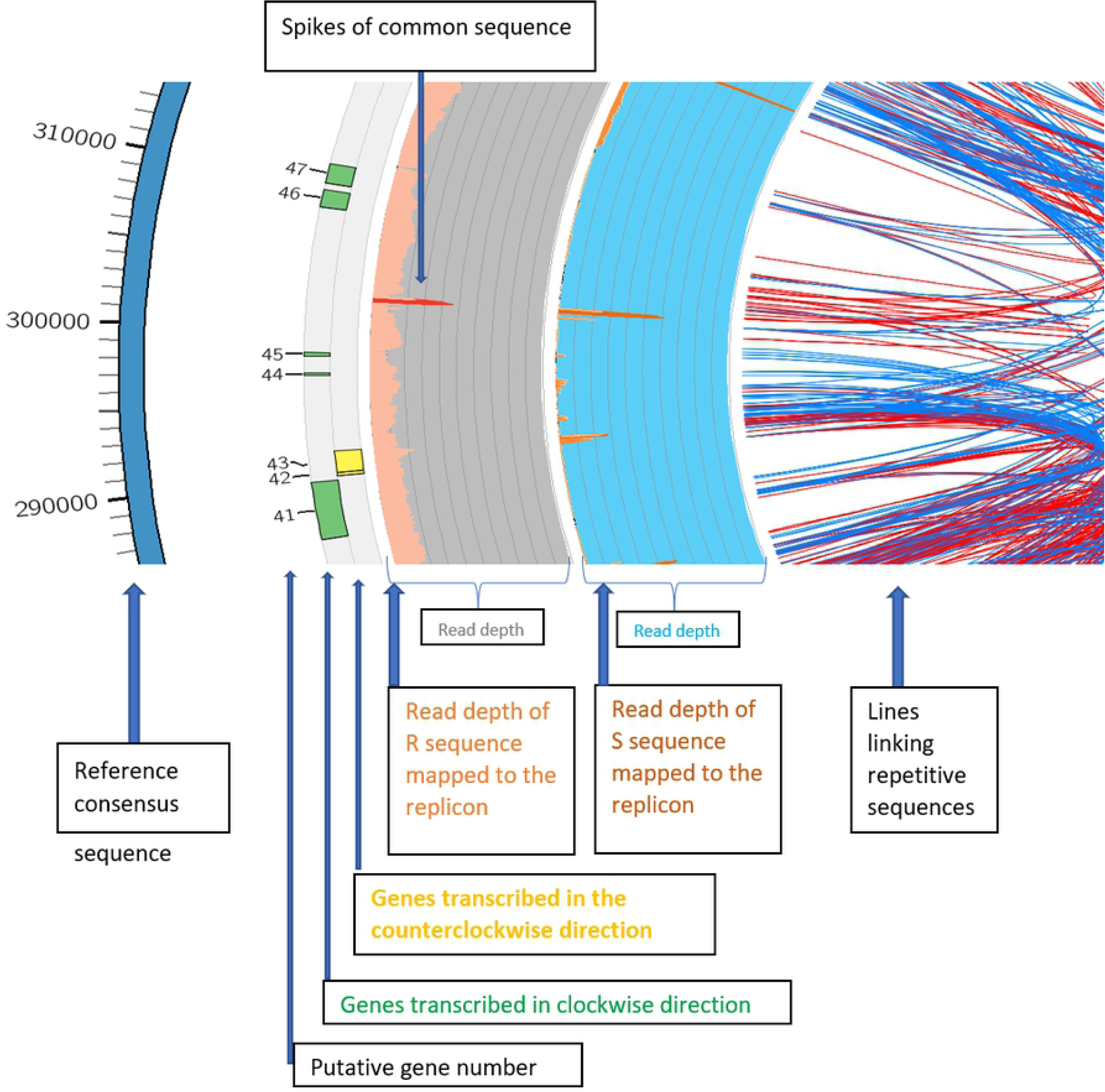
Enlargement of a segment of Fig. 1A. 300 kb showing a red/orange-colored spike representing regions of high copy numbers relative to the rest of the genome that has been acquired by the *EPSPS* replicon. The light pink band along the outer edge of the gray track is the read depth for a GR plant and indicates a read depth about 2000 whereas a similar continuous track is not present from sensitive plants.

The central core of Fig. 1A shows red and blue links of repetitive sequences. The *EPSPS* gene was surrounded by dense concentrations of tandem repeats (indicated by the red line links) which terminate in putative gene sequences that were not found clustered in S plants (Fig. 1A). The tandem repeats flanked the *EPSPS* gene, but largely excluded the inverted repeats (indicated with blue links in the center). In the GS WGS sequences, these repeat regions are not depicted as ‘spikes’ in Fig. 1, in comparison to other regions of the eccDNA replicon, suggesting they are lower in abundance in the GS genome.

WGS sequencing was conducted on GR biotypes from five states (AZ, GA, KS, DE and MD) and GS biotypes from three states (AZ, GA, KS). An average of 58 million read pairs per sample (approximately 36X genome coverage based on 410 Mbp haploid genome size of the glyphosate sensitive genome) were collected. Read pairs were mapped to the *EPSPS* replicon consensus sequence to assess representation, contiguity, and variation between the GR and GS biotypes (Fig. 2A, 2B and Supplemental Table 1). Greatly contrasting alignment rates were found after filtering for concordant pair alignments and quality (Q>20, Table 1). Each GR biotype mapped at least 2.2 M read pairs, with greater than 92% aligning in proper pairs (Fig. 1A). Conversely, the WGS reads derived from GS biotypes had less than 200 k high quality aligned pairs, where 50% or less were in the proper orientation (Fig 2A and 2B). Average read mapping depths of GS biotype sequences to the replicon varied by a magnitude of 5. All GR biotypes had at least 1,000 reads mapped when averaged; while all GS biotypes had no more than 181 reads as an average depth (Supplemental Table 1). When Illumina data from GS biotypes was aligned to the GR replicon, there was a striking lack of sequence contiguity; furthermore, many regions of the GR replicon had no Illumina data that aligned to it suggesting those regions were missing from the sampled GS biotypes. Therefore, we conclude that the *EPSPS* replicon is not present in its entirety in the background of the *A. palmeri* genome and was somehow assembled from many pieces from distal parts of the genome.

**Figure 2A.**
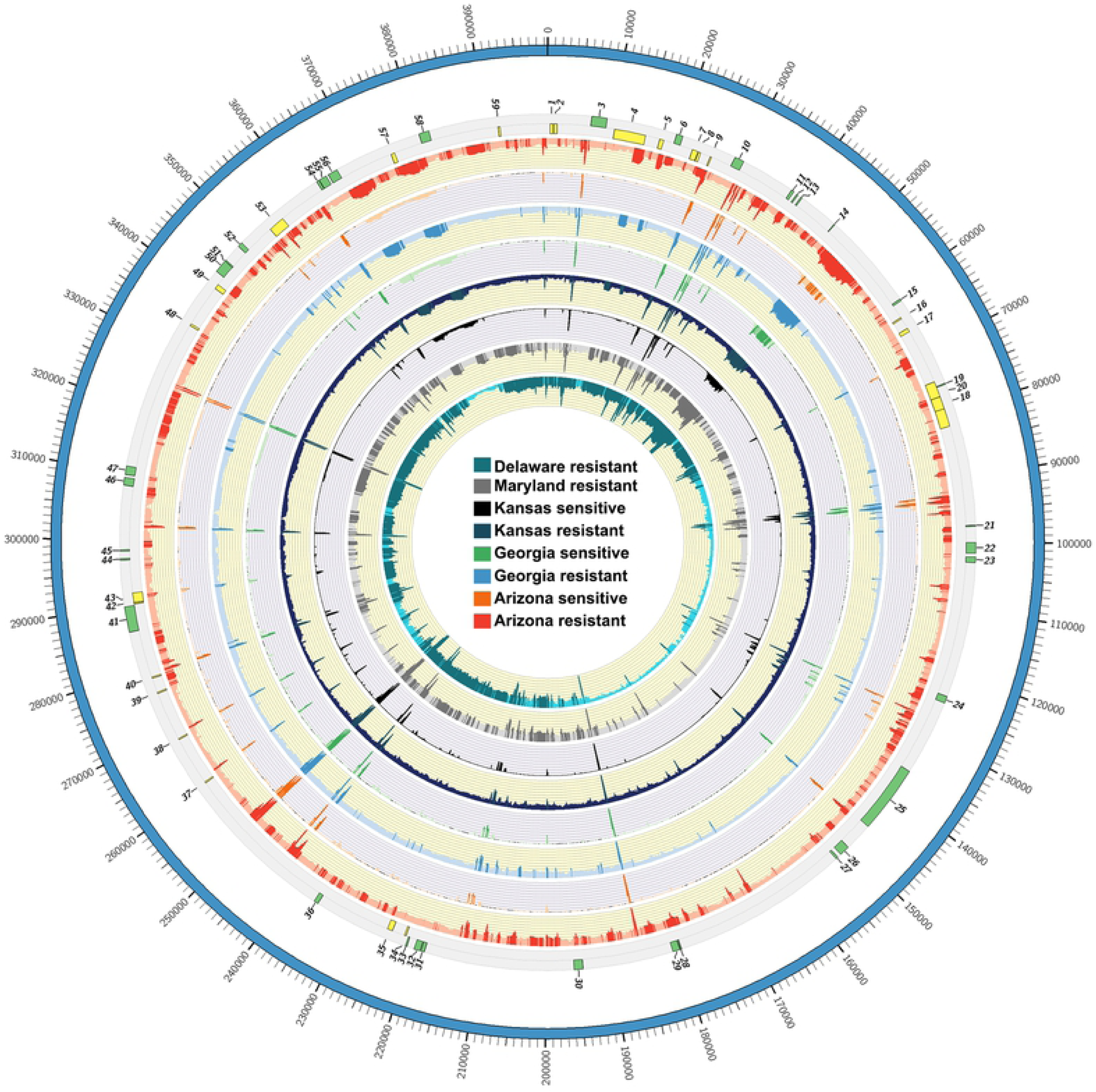
Mapping and alignment of WGS reads from R and S plants. (a) Mapping and alignment of WGS reads from GR and GS plants from different locations and populations from across the USA to the EPSPS replicon.

**Figure 2B.**
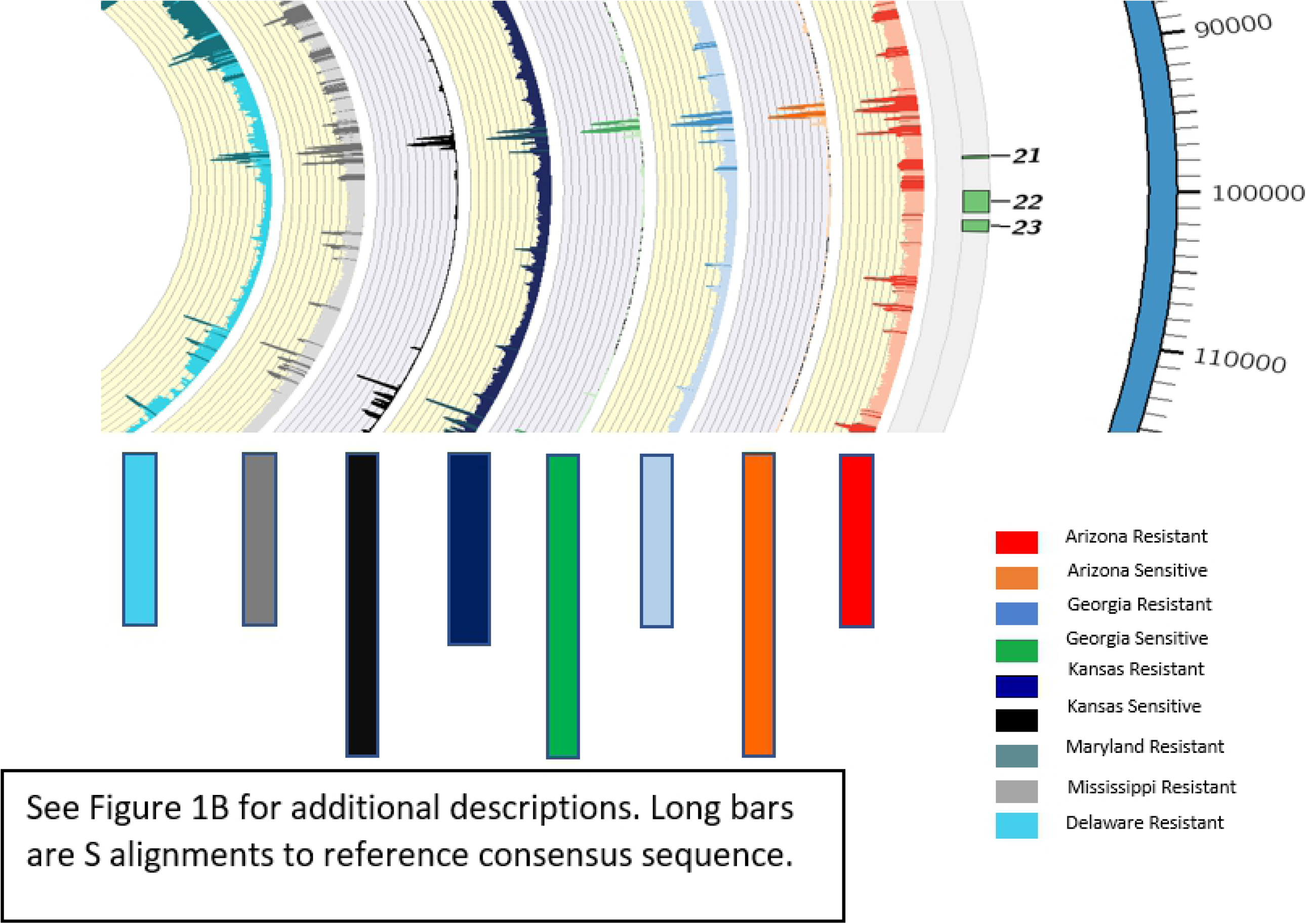
Enlarged segment of Fig. 2A. From left to right, short arrows point to color matched read depths of GR plants from Delaware (DE-R), Maryland (MD-R), Kansas (KS-R), Georgia (GA-R), and Arizona (AZ-R aligned to the Mississippi reference assembly and long arrows point to color matched read depth of GS plants from Kansas (KS-S), Georgia (GA_S) and Arizona (AZ_S) alignments.

The WGS reads from the three GS and five GR biotypes were aligned against the whole-genome assembly of a susceptible individual from the AZ-S population with the addition of the full-length replicon. These alignments were analyzed using CNVnator for the detection of other putative CNV events and/or replicons. CNVnator detected several putative duplications and deletions, most with relatively small changes in read depth (Supplementary Table 2). Some CNV events indicated extremely high amounts of increased read depth; however, besides the EPSPS replicon, no other copy number variant was detected that distinguished GR from GS biotypes, i.e. that was unique in all GR and not GS or *vice versa*. The read depth and relative copy number of all 59 genes in the GR replicon was homogenous, indicating that the replicon does not exist in multiple configurations either within an individual or across individuals from distant geographic regions. Some putative CNV events were biotype specific or regionally specific (i.e. KS, GA, AZ, MD, DE). These were generally few and did not show drastic deviation from background read depth. There were also several putative CNVs detected in all eight biotypes, indicating the genome was incomplete and did not accurately represent highly repetitive elements. (Supplemental Table 2).

The question of whether the *EPSPS* replicon evolved once and rapidly moved across the USA or evolved many times in different locations was addressed by assessing single nucleotide polymorphism (SNP) variation of the GR and GS biotypes from geographic distant locations from across the USA relative to the *EPSPS* replicon from Mississippi. Within the 399 kb sequence, a total of 5,079 SNPs and 738 Insertion/Deletions (INDELS) were identified among the variant sites that were genotyped (Table 2 and Supplemental Table 3). Consistently, the GS biotype samples contained approximately 4-fold more SNPs than the GR biotypes; while GR biotypes had very few private SNPs (<10), except for the Maryland sample which had only 277 private SNPs (Table 2).

**Table 2.** Genotyping statistics from WGS reads.

| State | Biotype | SNP Count | Indel Count | Private | Missing |
| --- | --- | --- | --- | --- | --- |
| Arizona | Resistant | 323 | 113 | 3 | 65 |
| Arizona | Sensitive | 2359 | 289 | 946 | 2162 |
| Delaware | Resistant | 495 | 166 | 10 | 48 |
| Georgia | Resistant | 465 | 133 | 4 | 65 |
| Georgia | Sensitive | 1858 | 133 | 413 | 2881 |
| Kansas | Resistant | 458 | 125 | 2 | 74 |
| Kansas | Sensitive | 2022 | 247 | 497 | 2203 |
| Maryland | Resistant | 945 | 179 | 277 | 380 |

Clustering of the samples based on SNPs revealed that the GR biotypes formed a distinct group from the GS biotypes (Figures 3A and 3B). The GR biotypes are on the same plane in the first principal component, and are only separated by the second principal component, with Maryland and Delaware appearing as slight outliers. This is also evident in Figure 3B, and both analyses indicate a high degree of relationship among GR types. The sequences from the different GR biotypes were also had similar sequence representation. Points along the replicon having greater read depths, which appear as spikes (Fig. 2A and 2B), also aligned perfectly with each other. Not only did these WGS datasets have highly conserved sequence, the contiguity among the genes and repetitive sequences along the length of the replicon was preserved in GR but not GS biotypes. Alignment of WGS reads from GS biotypes to the *EPSPS* replicon revealed gaps in the sequence in all the sensitive populations tested, and very low sequence coverage for the *EPSPS* replicon genes that are co-amplified in GR biotypes, as previously described [5].

**Figure 3.**
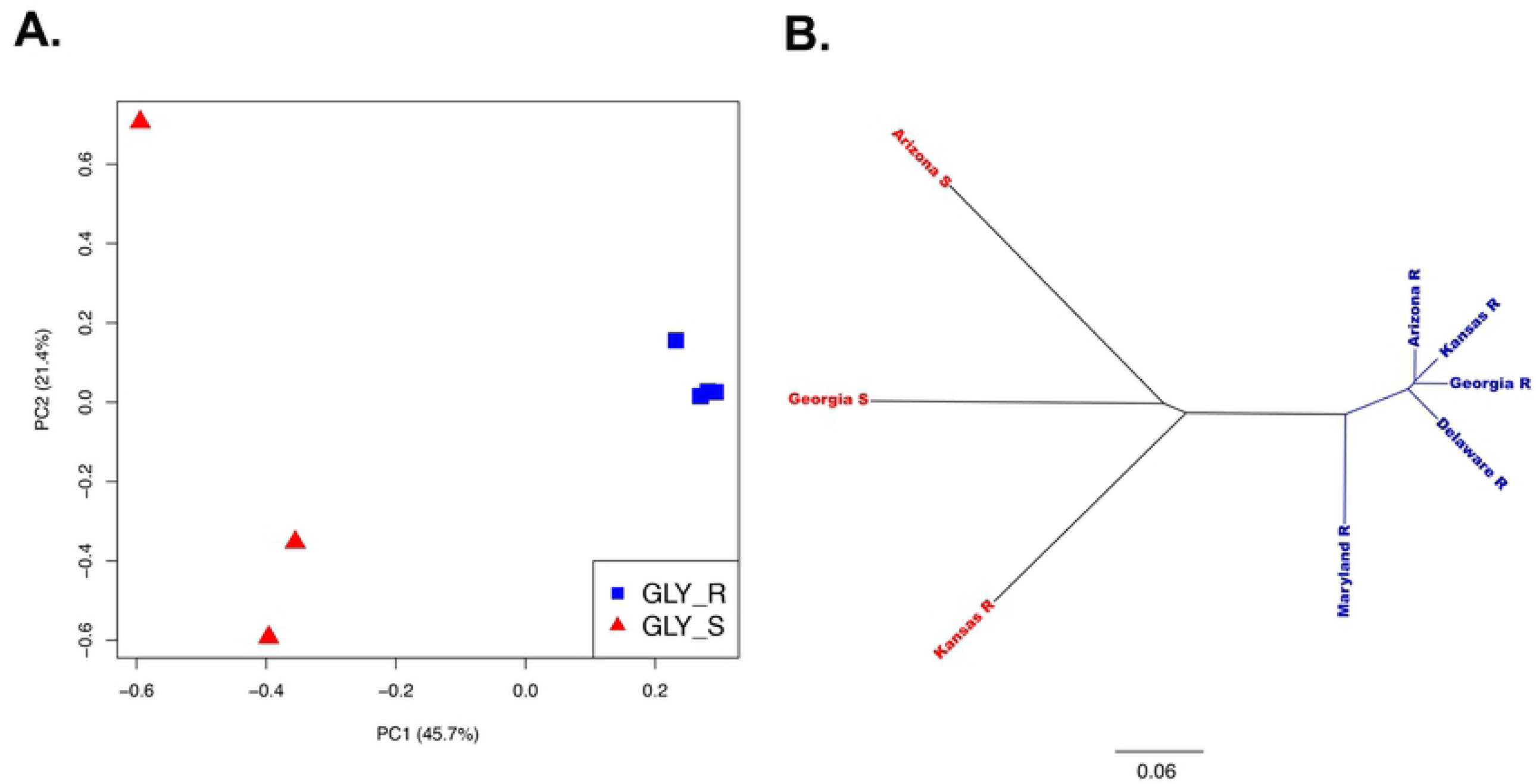
Principal component analysis of WGS reads from R and S plants. (A) Principal component analysis of WGS reads from GR and GS plants from different locations indicating the level of similarity among EPSPS replicons; (B) Cluster diagrams of the WGS reads from the GR and GS plants from different locations indicating the level of similarity among *EPSPS* replicons.

Although it cannot be stated that all cases of glyphosate resistance are due to the highly conserved *EPSPS* replicon, the GR samples sequenced here were nearly identical with consistent depth of reads and the exact size and position of components (spikes, genes) across six states with significant geographic separation between the GR biotypes when analyzed with the eccDNA replicon as a reference. Additionally, every gene in the replicon had an identical read depth, indicating that the entire replicon is present and at the same abundance (Supplemental Table 2). The read sequences mapped to the *EPSPS* consensus sequence with near perfect concordant and contiguous alignments with similar abundances of gene copies, repeat elements, and low-complexity sequence. The mapped reads compared across replicon sequences from different locations also contained very few polymorphisms (SNPs/INDELS) and read depths were consistent and greater than 1k, which provided further evidence that the replicons are significantly amplified in GR biotypes. Moreover, SNP diversity of the amplified genes has been reduced to singular homologous forms in the *EPSPS* replicon because of expansion by high copy numbers of the singular replicon sequence. Overall, there has been a reduction of genetic variation for the *EPSPS* which also points to the possibility that the replicon and replicon-based resistance appears to be a singular event. Hence, these results support the hypothesis that the replicon originated in one location, possibly from a single source, then rapidly spread across the United States through human or other natural dispersal means.

The replicon not only provides a unique heritable unit enabling adaption to herbicide stress, but also adds another 58 putative genes (aside from *EPSPS*) and gene products which could conceivably alter the genomic trajectory of this species. Glyphosate resistance has not negatively impacted *A. palmeri* fitness [8-10] and the additional 41 expressed genes may provide advantages in areas such as abiotic and biotic stress tolerance [11]. Replicon-based glyphosate resistance represents a natural, molecular solution constructed, modified and initiated by *A. palmeri*, a consequence that may inform future efforts in biomolecular engineering of genes that promote survival and genomic success.

The replicon can be found both integrated within the nuclear genome and tethered externally to chromosomes indicating the presence of bimodal forms of transmission of the replicon to daughter cells [14]. The mechanisms regulating distribution patterns of replicons in dividing cells is unknown, but may be determined by the number of available tethering sites as well as the number of copies of replicons present in the genome. Inheritance patterns are still poorly understood, but current data supports a non-mendelian inheritance pattern [9, 15-19] that may be driven by disproportionate replicon association in the genome during cell division. This may account for the copy number variations observed in *A. palmeri* populations and explains why F2 offspring from a resistant and susceptible test cross do not show the predicted 3:1 segregation of resistant:susceptible phenotype [15-19]. This also implies that the amplified *EPSPS* sequence and other genes within the replicon may not be genetically linked with any other trait in the genome and undergo independent assortment from the rest of the genome, which in turn, implies inheritance of the *EPSPS* resistance trait (i.e. the *EPSPS* replicon) does not extort the same amount of linkage drag [29, 30]. Once a resistant plant from a distant location crosses with a locally adapted susceptible plant, the resulting offspring can backcross with the local susceptible plants and purge any negative traits that may have been initially co-introduced with glyphosate resistance.

The origins of the replicon, from its initial stages to its present-day configuration are unknown, but likely occurred in a short evolutionary window. The replicon may have been formed through circularization hotspots [31], or from genome shuffling, reorganization, and recombination events within the genome with influence from mobile genetic elements [32]. Circularization hotspots, while again unknown, may be regions of repetitive sequences capable of intra-molecular recombination. Such regions are common in the replicon. Parts of the *EPSPS* replicon as partial or intermediary forms, or its building blocks, have not been found to be contiguous or in similar abundance in GS plants. The conditions responsible for the selection pressure under which the replicon was initially formed are also unknown. Elucidating the compounded evolutionary events leading to its development, such as chromatin reorganization/genome reshuffling, and evidence for segmental duplications and non-homologous end joining events, would improve our understanding of how such a structure was generated. The added genomic content, whereas it does result in an imbalance in gene dosage, does not appear to rise to a case of aneuploidy because the replicon is much smaller than the entire chromosome, lacks a centromere, and it exhibits nonmendelian genetics. Finally, due to the conservation of the nucleotide content and structure of the *EPSPS* replicon in the geographically diverse populations used in this study, the results strongly suggest that the events that gave rise to this form of glyphosate resistance likely only occurred once.

## Author contributions

Conceived and designed the experiments: WTM, CAS, ELP. Performed the experiments: WTM, CAS. Analyzed the data: WTM, CAS, ELP. Contributed reagents/materials/analysis tools: WTM, CAS. Wrote the paper: WTM, CAS, ELP.

## Supporting information captions

**Supplemental Table 1. Read depths for each wgs dataset aligned to the eccDNA replicon.** Number of whole genome shotgun reads that align to each nucleotide of the eccDNA replicon reference assembly. Average read depths are in parenthesis.

**Supplemental Table 2. CNVnator Read Depths of putative CNVs.** This table presents relative read depths as determined by CNVnator for each gene in the *Amaranthus palmeri* and eccDNA replicon reference assemblies. A gene is considered duplicated when relative read depths are greater than 2.

**Supplemental Table 3. SNPs by nucleotide position in the cassette.** The REF column is the eccDNA replicon reference allele. X/X indicates the plant is homozygous for the reference. X/YYY indicates there is an insertion../. Indicates the genotype is missing (no reads at this position).

## References

1. Culpepper AS, Grey TL, Vencill WK, Kichler JM, Webster TM, Brown SM, et al. Glyphosate-resistant Palmer amaranth confirmed in Georgia. Weed Sci. 2006;54: 620–626.

2. Heap, I. The International Survey of Herbicide Resistant Weeds. Accessed: Friday March 24, 2020. Available from: http://www.weedscience.org.

3. Gaines TA, Zhang W, Wang D, Bukun B, Chisholm ST, Shaner DL, et al. Gene amplification confers glyphosate resistance in Amaranthus palmeri. Proc Natl Acad Sci USA. 2010;107: 1029–1034.

4. Gaines TA, Wright AA, Molin WT, Lorentz L, Riggins CW, Tranel PJ, et al. Identification of genetic elements associated with EPSPS gene amplification. PLOS ONE. 2013;8: e65819. Available from: http://dx.doi.org/10.1371/journal.pone.0065819.

5. Molin WT, Wright AA, Lawton-Rauh A, Saski, CS. The unique genomic landscape surrounding the EPSPS gene in glyphosate resistant Amaranthus palmeri: A repetitive path to resistance. BMC Genomics. 2017;18: 91. Available from: doi: 10.1186/s12864-016-3336-4.

6. Molin WT, Wright AA, VanGessel MJ, McCloskey WB, Jugulam M, Hoagland RE. Survey of the genomic landscape surrounding the 5-enolpyruvylshikimate-3-phosphate synthase (EPSPS) gene in glyphosate-resistant Amaranthus palmeri from geographically distant populations in the USA. Pest Manag Sci. 2017; Available from: doi: 10.1002/ps.4659.

7. Chahal PS, Varanasi VK, Jugulam M, Jhala AJ. Glyphosate-Resistant Palmer Amaranth (Amaranthus palmeri) in Nebraska: Confirmation, EPSPS Gene Amplification, and Response to POST Corn and Soybean Herbicides. Weed Technol. 2017; 31: 80–93.

8. Giacomini D, Westra, P, Ward SM. Impact of genetic background in fitness cost studies: An Example from glyphosate-resistant Palmer Amaranth. Weed Sci. 2014; 62:29–37.

9. Vila-Aiub MM, Goh SS, Gaines TA, Han H, Busi R, Yu Q, Powles SB. No fitness cost of glyphosate resistance endowed by massive EPSPS gene amplification in Amaranthus palmeri. Planta 2014; doi 10.1007/s00425-013-2022-x

10. Vila-Aiub MM, Qin Yu Q, Powles SB. Do plants pay a fitness cost to be resistant to glyphosate? New Phytologist 2019, 223: 532–547. doi: 10.1111/nph.15733

11. Molin WT, Yaguchi A, Blenner M, Saski CA. The eccDNA Replicon: A heritable, extra-nuclear vehicle that enables gene amplification and glyphosate resistance in Amaranthus palmeri. The Plant Cell. 2020. DOI: https://10.1105/tpc.20.00099. First Published on April 23, 2020.

12. Hirochika H, Otsuk, H. Extrachromosomal circular forms of the tobacco retrotransposons Tto1. Gene. 1995;165: 229–232.

13. Lanciano S, Carpentier M-C, Llauro C, Jobet E, Robakowska-Hyzorek D, Lasserre E, et al. Sequencing the extrachromosomal circular mobilome reveals retrotransposon activity in plants. PLoS Genet. 2017; 13(2): e1006630. doi: 10.1371/journal.pgen.1006630.

14. Koo D-H, Molin WT, Saski CA, Jiang J, Putta K, Jugulam M, et al. Extra-chromosomal circular DNA-based amplification and transmission of herbicide resistance in crop weed Amaranthus palmeri. Proc Nat Acad Sci USA. 2018; http://www.pnas.org/cgi/doi/10.1073/pnas.1719354115.

15. Gaines, T. A., D. L. Shaner, S. M. Ward, J. E. Leach, C. Preston, and P. Westra 2011. Mechanism of resistance of evolved glyphosate resistant Palmer amaranth (Amaranthus palmeri). Journal of Agricultural and Food Chemistry 59:5886–5889.

16. Chandi A, Milla-Lewis SR, Giacomini D, Westra P, Preston C, Jordan DL, York AC, Burton JD, Whitaker JR. Inheritance of evolved glyphosate resistance in a North Carolina Palmer amaranth (Amaranthus palmeri) biotype. Int J Agron. 2012; Article ID: 176108. doi: 10.1155/2012/176108.

17. Giacomini DA, Westra P, Ward SM. Variable Inheritance of Amplified EPSPS Gene Copies in Glyphosate-Resistant Palmer Amaranth (Amaranthus palmeri). Weed Sci. 2019; 67: 176–182. Available from doi: 10.1017/wsc.2018.65

18. Patterson EL, Pettinga DJ, Ravet K, Neve P, Gaines TA. Glyphosate Resistance and EPSPS Gene Duplication: Convergent Evolution in Multiple Plant Species. J.Hered. 2018; 117–125; doi: 10.1093/jhered/esx087

19. Sammons RD, Gaines TA. Glyphosate resistance: state of knowledge. Pest Manag. Sci. 2014, (wileyonlinelibrary.com) DOI 10.1002/ps.3743

20. Küpper A, Manmathan HK, Giacomini D, Patterson EL, McCloskey WB, Gaines TA. Population genetic structure in glyphosate-resistant and -susceptible Palmer amaranth (Amaranthus palmeri) populations using genotyping-by-sequencing (GBS). Frontiers in Plant Sci. 2018; 9:29. Available from: https://doi.org/10.3389/fpls.2018.00029.

21. Bolger AM, Lohse M, Usadel B. Trimmomatic: a flexible trimmer for Illumina sequence data. Bioinformatics. 2014;30: 2114–2120.

22. Langmead B, Salzberg SL. Fast gapped-read alignment with Bowtie 2. Nat. Methods. 2012;9: 357–359.

23. Li H, Handsaker B, Wysoker A, Fennell T, Ruan J, Homer N, et al. 1000 Genome Project Data Processing Subgroup. The Sequence alignment/map format and SAMtools. Bioinformatics. 2009;25: 2078–2079.

24. Danecek P, Auton A, Abecasis G, Albers CA, Banks E, DePristo MA, et al. 1000 Genomes Project Analysis Group, The variant call format and VCFtools. Bioinformatics. 2011;27: 2156–2158.

25. Krzywinski M, Schein J, Birol I, Connors J, Gascoyne R, Horsman D, et al. Circos: an information aesthetic for comparative genomics. Genome Res. 2009;19: 1639-1645.23.

26. Abyzov A, Urban AE, Snyder M, Gerstein M. CNVnator: an approach to discover, genotype, and characterize typical and atypical CNVs from family and population genome sequencing. Genome Res. 2011;21: 974–84.

27. Mills RE, Walter K, Stewart C, Handsaker RE, Chen K, Alkan C, et al. Mapping copy number variation by population-scale genome sequencing. Nature. 2011; Feb;470(7332): 59.

28. Quinlan AR, Hall IM. BEDTools: a flexible suite of utilities for comparing genomic features. Bioinformatics. 2010;26: 841–842.

29. Brown AHD, Lawrence GJ, Jenkin M, Douglass J, Gregory E. Linkage drag in backcross breeding in barley. J Hered. 1989;80: 234–239.

30. Zeven AC, Knott DR, Johnson R. Investigation of linkage drag in near isogenic lines of wheat by testing for seedling reaction to races of stem rust, leaf rust and yellow rust. Euphytica. 1983;32: 319–327.

31. Moller HD, Mohiyuddin M, Prada-Luengo I, et al. Circular DNA elements of chromosomal origin are common inhealthy human somatic tissue. Nature Commun. 2018; 9(1) doi: 10.1038/s41467-018-03369-8

32. Moller HD, Larsen CE, Parsons L, Hansen AJ, Regenberg B, Mourier T. Formation of extrachromosomal circular DNA from long terminal repeats of retrotransposons in *Saccharomyces cerevisiae*. Genes, Genes, Genetics 2016;6: 453–462. https://doi.org/10.1534/g3.115.025858

